# Adapting protein language models for rapid DTI prediction

**DOI:** 10.1101/2022.11.03.515084

**Authors:** Samuel Sledzieski, Rohit Singh, Lenore Cowen, Bonnie Berger

## Abstract

We consider the problem of sequence-based drug-target interaction (DTI) prediction, showing that a straightforward deep learning architecture that leverages pre-trained protein language models (PLMs) for protein embedding outperforms state of the art approaches, achieving higher accuracy, expanded generalizability, and an order of magnitude faster training. PLM embeddings are found to contain general information that is especially useful in few-shot (small training data set) and zero-shot instances (unseen proteins or drugs). Additionally, the PLM embeddings can be augmented with features tuned by task-specific pre-training, and we find that these task-specific features are more informative than baseline PLM features. We anticipate such transfer learning approaches will facilitate rapid prototyping of DTI models, especially in low-N scenarios.

## 1 Introduction

Predicting drug-target interaction (DTI), a critically important problem in drug discovery, should ideally be informed by protein and drug structures. However, even if all protein structures were available (say, by AlphaFold2 prediction [14]), the computational expense of docking is prohibitive for large-scale DTI screening, suggesting that sequence-based prediction of DTIs will remain important.

Accordingly, in this paper we consider the computational prediction of DTIs when the inputs are a) a molecular description of the drug (such as the SMILES string [1]) and b) the amino acid sequence of the protein target. Many methods have been proposed to address the DTI problem in this formulation [2], with state-of-the-art approaches relying on deep learning architectures that build protein representations using sequence models like convolutional neural networks [15] and, more recently, transformers [11]. As the focus of this work is optimal protein representations for DTI prediction, we fix the drug molecular representation here to the commonly used Morgan fingerprints [17], noting that recently introduced alternative representations may further increase performance [12, 13].

The key claim of this work is that a pre-trained protein representation from protein language models can offer state-of-the-art performance with a relatively straightforward model architecture, even on tasks that have already been the focus of dedicated deep learning model design. Pre-training is especially impactful when the training set is small or unbalanced or if the test set contains hitherto unseen proteins or drugs. We emphasize here the importance of matching the pre-trained representation to the task. As we show in a feature attribution analysis, augmenting language models with additional training on protein-protein interaction (PPI) prediction yields features which are more informative for DTI prediction than those from baseline embeddings that were pre-trained only on protein sequences. Thus, our work constitutes a concrete demonstration of the power of a well-designed transfer learning approach that adapts foundation models for a specific task [4, 7].

## 2 Method

Given a SMILES string and protein sequence, our predictive framework consists of the following three steps (Figure 1):

**Figure 1:**
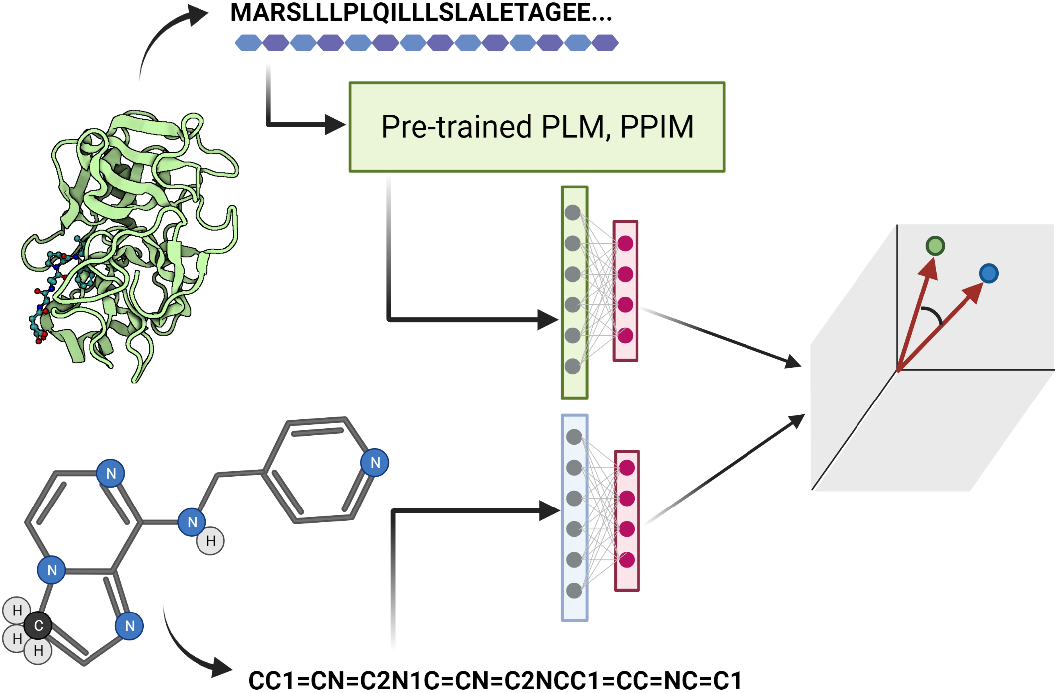
Our prediction framework leverages protein sequence representations learned by pre-trained protein langauage models (PLM) and protein-protein interaction prediction models (PPIM) to make accurate drug-target interaction predictions. Pre-trained features generalize well to unseen proteins, allowing for zero- and few-shot learning of DTIs.

**Figure 2:**
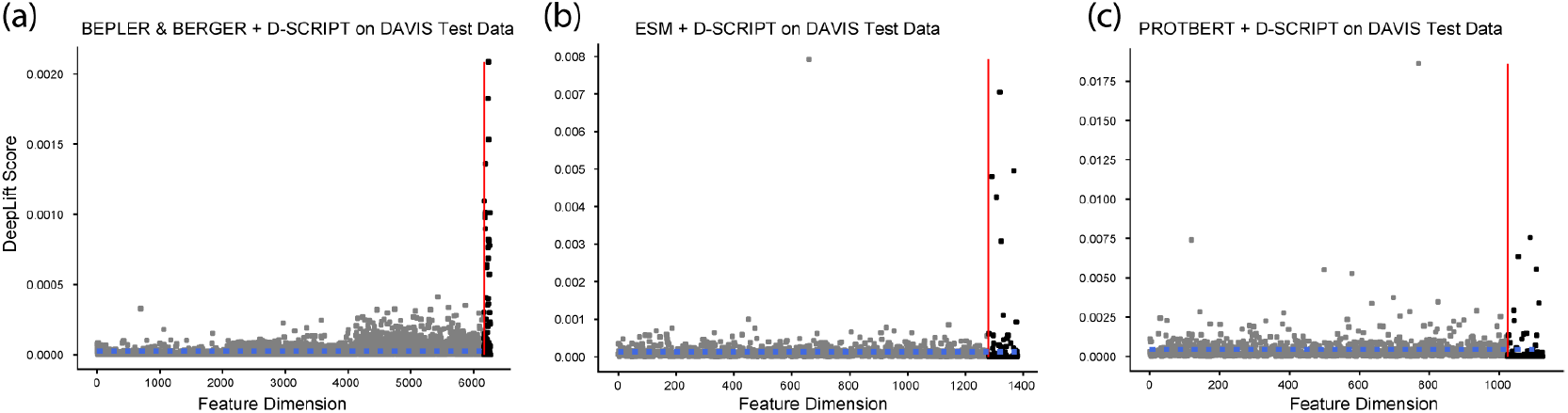
DeepLift feature attributions for Bepler & Berger **(a)**, ESM **(b)**, and ProtBert **(c)** embedding dimensions (gray) and the respective D-SCRIPT embedding dimensions (black) of the combined featurizations from Section 3.2. D-SCRIPT features have an outsize contribution to the overall model prediction relative to the PLM features alone. The dashed blue line indicates the mean feature attribution.

1. Featurize the drug and protein using pre-trained models or algorithms.
2. Transform both the drug and protein into a shared latent space.
3. Output the DTI prediction based on the drug–protein distance in the latent space.

### Molecular featurization

We featurize the drug molecule by its Morgan fingerprint [17], an encoding of the SMILES string of the molecular graph as a fixed-dimension embedding 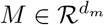 (we chose *d*_*m*_ = 2048) by considering the local neighborhood around each atom. The utility of the Morgan fingerprint for small molecule representation has been demonstrated in [20].

### Pre-trained protein featurization

We generate protein features using pre-trained protein language models (PLM): These models generate a protein embedding *E*^+^ ∈ ℛ^*n*×*d*^ for a protein of length *n*, which is then pooled along the length of the protein resulting in a vector *E* ∈ ℛ^*d*^. Specifically, we investigate the PLMs from Bepler & Berger [3], ESM [19], and ProtBert [6], with default dimensions *d* = 6165, 1280, 1024 respectively.

Additionally, we evaluate the output of the projection module of a D-SCRIPT PPI prediction model trained on human PPIs, using each language model as input embeddings [22]. Details of this featurization (*d* = 100) can be found in Appendix A.1 We emphasize that the language and projection models are used exclusively to generate input features– their weights are kept unchanged and are not updated during DTI training.

### Transformation into a shared latent space

Given small molecule embedding 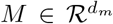 and protein embedding *E* ∈ ℛ^*d*^, we transform them separately into *M* ^*^, *E*^*^ ∈ ℛ^*h*^ using fully-connected multi-layer perceptrons with a ReLU activation. Given the latent embeddings *M* ^*^, *E*^*^, we compute the probability of a drug-target interaction 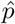 as the cosine similarity between the embedding vectors.

### Training and Implementation

The loss was calculated using the binary cross entropy between the true labels *y* and the predicted interaction probabilities 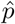. Model weights were updated with error back-propagation using the Adam optimizer with learning rate 10^−4^ over 50 epochs, with a batch size of 16. We used a latent dimension size of 1024 (results were robust to variations in latent dimension size). We implemented this framework in PyTorch version 1.9.

## 3 Results

### Data Sets

To evaluate the predictive accuracy of our framework, we use three different DTI benchmark data sets. Two data sets, **DAVIS** [5] and **BindingDB** [16], consist of pairs of drugs and targets with experimentally determined dissociation constants (*K*_*d*_). Following [11], we treat pairs with *K*_*d*_ *<* 30 as positive DTIs, while larger *K*_*d*_ values are negative. The third data set, ChG-Miner from **BIOSNAP** [24], consists of only positive DTIs. The DAVIS data set represents a few-shot learning setting: it contains only 2,086 training interactions, compared to 12,668 for BindingDB and 19,238 for BIOSNAP. The rest of the data preparation follows [11]. We create negative DTIs by randomly sampling an equal number of protein-drug pairs, with the expectation that a random pair is unlikely to be positively interacting. The data sets are split into 70% for training, 10% for validation, and the remaining 20% for testing. Training data is artificially sub-sampled to have an equal number of positive and negative interactions, while validation and test data is left at the natural ratio. Full specification of the data, including number of unique drugs, proteins, and positive/negative edges can be found in Table A1.

### Experiment Design

For each data set, we evaluate the predictive performance of the pre-trained **Bepler & Berger, ESM**, and **ProtBert** model embeddings. Additionally, we compare with **MolTrans** [11], **GNN-CPI** [23], and **DeepConv-DTI** [15], which have been shown to achieve state-of-the-art performance on DTI prediction, and specifically on these benchmark data sets. During training, we monitor the AUPR, AUROC, and F1 metrics on the validation set, and store the model with the highest AUPR, which is then evaluated on the held-out testing set. Each training is run with 5 random initializations, and we report the mean and standard deviation of each metric.

#### 3.1 Improved DTI prediction

We demonstrate that co-embedding pre-trained protein language model features with small molecule features achieves state-of-the-art performance on all three benchmark data sets. Across all three data sets the three PLM models perform similarly, which is consistent with prior work which shows that there is often ambiguity as to which PLM is best suited to a given task [8, 9]. However, the three PLM models consistently outperform the non-PLM models, with especially large improvement coming on the most challenging DAVIS set where very little training data is available and evaluation is unbalanced. DeepConv-DTI achieves the best AUROC on the unbalanced BindingDB, but the PLM models have higher AUPR, which is the more representative metric for unbalanced data sets.

#### 3.2 Feature attribution reveals information gain from tuning on PPI

We additionally investigated training DTI models using protein language model embeddings augmented with features from a D-SCRIPT model pre-trained on human PPIs (see Appendix A.1 for details). While the top-line performance of the augmented models are similar to the base models (Table A3), an attribution study using DeepLift [21] shows that the new D-SCRIPT-derived features are disproportionately represented in the set of highly important features. This suggests that tuning on a related task refines the representations from the general protein language models to ones more suited for the specific task, as in [7]. This explanation is supported by the fact that the 100-dimensional D-SCRIPT features alone achieved only slightly decreased performance on the DTI task compared to PLM-based models with 10-50x as many parameters (Table A3).

#### 3.3 Zero-shot learning with pre-trained protein embeddings

In [22], Sledzieski et al. demonstrate that language models enable D-SCRIPT to generalize especially well to out-of-species PPIs. Here, we show that generalization extends to DTIs for proteins which are unseen in the training set. The Bepler & Berger | BB-D-SCRIPT featurization outperforms MolTrans in predictive performance on a variation of the BIOSNAP data where 20% of proteins and all corresponding interactions were removed as a test set. The outperformance over MolTrans is not as stark in the unseen drugs domain, possibly because the informational advantage of pre-training disproportionately benefits the protein representations (Table 2). The performance of other PLM models was similar to the one shown (Table A3).

**Table 1:** PLM-based models (Bepler & Berger, ESM, ProtBert) out-perform non-PLM models (MolTrans, GNN-CPI, DeepConv-DTI) over 5 random initializations. Training pairs are positive + negative, with an equal number of each. The column “Balanced?” refers to validation and test sets. Metrics for models with † are from [11]

| Benchmark | Training Edges | Balanced? | Model | AUPR | AUROC | F1 |
| --- | --- | --- | --- | --- | --- | --- |
| BIOSNAP | 19238 | Yes | Bepler & Berger | **0.914 ± 0.003** | **0.898 ± 0.004** | **0.838 ± 0.004** |
|  |  |  | ESM | 0.898 ± 0.004 | 0.876 ± 0.005 | 0.817 ± 0.008 |
|  |  |  | ProtBert | 0.895 ± 0.004 | 0.873 ± 0.004 | 0.811 ± 0.004 |
|  |  |  | MolTrans | 0.885 ± 0.005 | 0.876 ± 0.007 | 0.806 ± 0.006 |
|  |  |  | GNN-CPI† | 0.890 ± 0.004 | 0.879 ± 0.007 | — |
|  |  |  | DeepConv-DTI† | 0.889 ± 0.005 | 0.883 ± 0.002 | — |
| BindingDB | 12668 | No | Bepler & Berger | 0.618 ± 0.009 | 0.862 ± 0.006 | 0.625 ± 0.010 |
|  |  |  | ESM | 0.638 ± 0.005 | 0.881 ± 0.002 | **0.637 ± 0.003** |
|  |  |  | ProtBert | **0.652 ± 0.005** | 0.876 ± 0.007 | 0.636 ± 0.006 |
|  |  |  | MolTrans | 0.598 ± 0.013 | **0.898 ± 0.009** | 0.593 ± 0.015 |
|  |  |  | GNN-CPI† | 0.578 ± 0.015 | 0.900 ± 0.004 | — |
|  |  |  | DeepConv-DTI† | 0.611 ± 0.015 | 0.908 ± 0.004 | — |
| DAVIS | 2086 | No | Bepler & Berger | 0.463 ± 0.013 | 0.907 ± 0.005 | 0.523 ± 0.012 |
|  |  |  | ESM | 0.479 ± 0.008 | 0.916 ± 0.004 | 0.544 ± 0.008 |
|  |  |  | ProtBert | **0.511 ± 0.012** | **0.917 ± 0.003** | **0.546 ± 0.006** |
|  |  |  | MolTrans | 0.335 ± 0.017 | 0.889 ± 0.007 | 0.420 ± 0.012 |
|  |  |  | GNN-CPI† | 0.269 ± 0.020 | 0.840 ± 0.012 | — |
|  |  |  | DeepConv-DTI† | 0.299 ± 0.039 | 0.884 ± 0.008 | — |

**Table 2:** Pre-trained embeddings generalize DTI prediction to proteins not seen in training

| Benchmark | Model | AUPR | AUROC | F1 |
| --- | --- | --- | --- | --- |
| Unseen proteins | Bepler & Berger \| BB-D-SCRIPT | **0.875** | **0.868** | **0.793** |
|  | MolTrans | 0.657 | 0.660 | 0.664 |
| Unseen drugs | Bepler & Berger \| BB-D-SCRIPT | **0.882** | **0.858** | **0.779** |
|  | MolTrans | 0.858 | 0.832 | 0.761 |

#### 3.4 Pre-training enables an order of magnitude faster optimization

One of the benefits of using a hierarchy of pre-trained models is that computation times are amortized over the lifespan of downstream applications. Pre-trained models incur an up-front computational cost, but can then be re-used for multiple inference tasks with straightforward architectures. Our framework allows for training DTI models up to an order of magnitude faster than an end-to-end method. Inference of DTIs is also faster—here anywhere from a 2x to 5x speedup (Table A2).

## 4 Discussion

Previous work has recognized the value of meaningful drug molecule representations for DTI prediction [10, 18], but relatively little work has focused on the target protein representation. Here, we show that pre-trained embeddings from protein language models, combined with simple molecular features, not only achieve state-of-the-art performance for the DTI prediction task but also enable substantially better accuracy in the few-shot (DAVIS data set) or zero-shot (unseen proteins) learning settings. Also, features learned from pre-training on the related PPI prediction task can provide additional information beyond general protein language models. This approach enables generalization to unseen proteins as well as fast model training and inference. This is particularly valuable for drug re-purposing and iterative screening where large compound libraries are evaluated against hitherto-uncharacterized proteins from pathways implicated in a disease of interest. Our framework may enable more accurate transfer of DTI from the model organisms on which drugs are initially tested to their eventual use in human patients. This work demonstrates the previously unexplored value of language models in the DTI prediction domain, the additional information unlocked by pre-training on related tasks (PPI prediction), and the power of iterative adaption for transfer learning.

## Acknowledgements

SS was supported by the National Science Foundation Graduate Research Fellowship under Grant No. 1745302. RS and BB were supported by the NIH grant R35GM141861. LC was supported by NSF OAC-1939263 and CCF-1934553

## A Appendix

**Table A1:** Full specification of benchmark data sets. Number of pairs are shown as (positive/negative).

| Data Set | Unique Drugs | Unique Proteins | Training Pairs | Validation Pairs | Test Pairs |
| --- | --- | --- | --- | --- | --- |
| BIOSNAP | 4510 | 2181 | 9619/9619 | 1374/1374 | 2748/2748 |
| BindingDB | 10665 | 1413 | 6334/6334 | 927/5717 | 1905/11384 |
| DAVIS | 68 | 379 | 1043/1043 | 160/2846 | 303/5708 |

**Table A2:** Comparison of training and inference times of several models (mean seconds over 5 runs). Training and inference time for PLM models were all similar.

|  |  | Wall Clock | Training (per epoch) | Inference |
| --- | --- | --- | --- | --- |
| BindingDB | BB-D-SCRIPT | **848** | **4.65** | **42.46** |
|  | Bepler & Berger | 866 | 6.52 | 43.27 |
|  | MolTrans | 9874 | 142.96 | 222.17 |
| BIOSNAP | BB-D-SCRIPT | 496 | **5.35** | 18.12 |
|  | Bepler & Berger | **487** | 5.94 | **14.29** |
|  | MolTrans | 6424 | 116.21 | 31.64 |
| DAVIS | BB-D-SCRIPT | 231 | **0.57** | 10.19 |
|  | Bepler & Berger | **216** | 0.98 | **8.94** |
|  | MolTrans | 1417 | 15.38 | 35.06 |

### A.1 D-SCRIPT projections as protein features for DTI

In the original D-SCRIPT paper, protein sequence embeddings from the Bepler & Berger model are used as input features. Here, we additionally train D-SCRIPT models using ESM and ProtBert embeddings as features. These models are trained on the human cross-validation set from [22]. We refer to the output of the first projection module, which takes as input the *n* ×*d* protein representation and tunes it to an *n* × 100 representation, as [PLM]-D-SCRIPT, where PLM is either Bepler & Berger (BB), ESM, or ProtBert. As inputs for our DTI model, we explore concatenating these embeddings to the raw PLM embeddings (e.g. [ESM | ESM-D-DSCRIPT]) so each amino acid has *d* + 100 features, and using the D-SCRIPT projections on their own.

**Table A3:** Full results of all DTI models trained on all data sets. [PLM]-D-SCRIPT means a humantrained D-SCRIPT model where input features came from that protein-language model. [PLM] | [PLM]-D-SCRIPT means the D-SCRIPT embeddings concatenated to the raw embeddings.

| Benchmark | Training Edges | Balanced? | Model | AUPR | AUROC | F1 |
| --- | --- | --- | --- | --- | --- | --- |
| BIOSNAP | 19238 | Yes | [Bepler & Berger \| BB-D-SCRIPT] | 0.611 | 0.863 | 0.633 |
|  |  |  | BB-D-SCRIPT | 0.618 | 0.870 | 0.619 |
|  |  |  | Bepler & Berger | 0.618 | 0.862 | 0.625 |
|  |  |  | [ESM \| ESM-D-SCRIPT] | 0.641 | **0.881** | 0.638 |
|  |  |  | ESM-D-SCRIPT | 0.611 | 0.865 | 0.620 |
|  |  |  | ESM | 0.637 | **0.881** | 0.637 |
|  |  |  | [ProtBert \| ProtBert-D-SCRIPT] | **0.652** | 0.878 | **0.641** |
|  |  |  | ProtBert-D-SCRIPT | 0.624 | 0.863 | 0.623 |
|  |  |  | ProtBert | 0.652 | 0.876 | 0.636 |
| BindingDB | 12668 | No | [Bepler & Berger \| BB-D-SCRIPT] | 0.909 | 0.893 | 0.836 |
|  |  |  | BB-D-SCRIPT | 0.911 | 0.897 | 0.831 |
|  |  |  | Bepler & Berger | **0.914** | **0.898** | **0.838** |
|  |  |  | [ESM \| ESM-D-SCRIPT] | 0.901 | 0.881 | 0.823 |
|  |  |  | ESM-D-SCRIPT | 0.889 | 0.867 | 0.798 |
|  |  |  | ESM | 0.898 | 0.876 | 0.817 |
|  |  |  | [ProtBert \| ProtBert-D-SCRIPT] | 0.897 | 0.876 | 0.812 |
|  |  |  | ProtBert-D-SCRIPT | 0.885 | 0.866 | 0.796 |
|  |  |  | ProtBert | 0.895 | 0.873 | 0.811 |
| DAVIS | 2086 | No | [Bepler & Berger \| BB-D-SCRIPT] | 0.464 | 0.913 | 0.537 |
|  |  |  | BB-D-SCRIPT | 0.475 | 0.912 | 0.533 |
|  |  |  | Bepler & Berger | 0.463 | 0.907 | 0.523 |
|  |  |  | [ESM \| ESM-D-SCRIPT] | 0.495 | **0.920** | **0.547** |
|  |  |  | ESM-D-SCRIPT | 0.479 | 0.915 | 0.522 |
|  |  |  | ESM | 0.480 | 0.916 | 0.544 |
|  |  |  | [ProtBert \| ProtBert-D-SCRIPT] | **0.502** | 0.916 | 0.533 |
|  |  |  | ProtBert-D-SCRIPT | 0.462 | 0.903 | 0.522 |
|  |  |  | ProtBert | 0.511 | 0.917 | 0.546 |
| BIOSNAP Unseen Proteins |  |  | [Bepler & Berger \| BB-D-SCRIPT] | **0.875** | **0.868** | 0.793 |
|  |  |  | Bepler & Berger | **0.875** | **0.868** | **0.798** |
|  |  |  | [ESM \| ESM-D-SCRIPT] | 0.847 | 0.836 | 0.770 |
|  |  |  | ESM | 0.850 | 0.839 | 0.770 |
|  |  |  | [ProtBert \| ProtBert-D-SCRIPT] | 0.841 | 0.827 | 0.755 |
|  |  |  | ProtBert | 0.841 | 0.827 | 0.750 |
| BIOSNAP Unseen Drugs |  |  | [Bepler & Berger \| BB-D-SCRIPT] | 0.879 | 0.855 | 0.800 |
|  |  |  | Bepler & Berger | 0.881 | **0.857** | **0.802** |
|  |  |  | [ESM \| ESM-D-SCRIPT] | **0.882** | 0.853 | 0.793 |
|  |  |  | ESM | 0.876 | 0.850 | 0.796 |
|  |  |  | [ProtBert \| ProtBert-D-SCRIPT] | 0.877 | 0.851 | 0.791 |
|  |  |  | ProtBert | 0.876 | 0.848 | 0.785 |

## References

[1] E. Anderson, G. D. Veith, and D. Weininger. SMILES, a line notation and computerized interpreter for chemical structures. US Environmental Protection Agency, Environmental Research Laboratory, 1987.

[2] M. Bagherian, E. Sabeti, K. Wang, M. A. Sartor, Z. Nikolovska-Coleska, and K. Najarian. Machine learning approaches and databases for prediction of drug–target interaction: a survey paper. Briefings in Bioinformatics, page 23, 2021.

[3] T. Bepler and B. Berger. Learning protein sequence embeddings using information from structure. In 7th International Conference on Learning Representations, ICLR 2019, 2019.

[4] R. Bommasani, D. A. Hudson, E. Adeli, R. Altman, S. Arora, S. von Arx, M. S. Bernstein, J. Bohg, A. Bosselut, E. Brunskill, et al. On the opportunities and risks of foundation models. arXiv preprint arXiv:2108.07258, 2021.

[5] M. I. Davis, J. P. Hunt, S. Herrgard, P. Ciceri, L. M. Wodicka, G. Pallares, M. Hocker, D. K. Treiber, and P. P. Zarrinkar. Comprehensive analysis of kinase inhibitor selectivity. Nature biotechnology, 29(11):1046–1051, 2011.

[6] A. Elnaggar, M. Heinzinger, C. Dallago, G. Rihawi, Y. Wang, L. Jones, T. Gibbs, T. Feher, C. Angerer, M. Steinegger, et al. Prottrans: towards cracking the language of life’s code through self-supervised deep learning and high performance computing. arXiv preprint arXiv:2007.06225, 2020.

[7] S. Gururangan, A. Marasović, S. Swayamdipta, K. Lo, I. Beltagy, D. Downey, and N. A. Smith. Don’t stop pretraining: adapt language models to domains and tasks. arXiv preprint arXiv:2004.10964, 2020.

[8] B. L. Hie, K. K. Yang, and P. S. Kim. Evolutionary velocity with protein language models. bioRxiv, 2021.

[9] C. Hsu, H. Nisonoff, C. Fannjiang, and J. Listgarten. Combining evolutionary and assay-labelled data for protein fitness prediction. bioRxiv, 2021.

[10] K. Huang, T. Fu, L. M. Glass, M. Zitnik, C. Xiao, and J. Sun. DeepPurpose: a deep learning library for drug–target interaction prediction. Bioinformatics, 36(22-23):5545–5547, Apr. 2021.

[11] K. Huang, C. Xiao, L. M. Glass, and J. Sun. MolTrans: Molecular Interaction Transformer for drug–target interaction prediction. Bioinformatics, 37(6):830–836, May 2021.

[12] W. Jin, R. Barzilay, and T. Jaakkola. Junction tree variational autoencoder for molecular graph generation. In International Conference on Machine Learning, pages 2323–2332. PMLR, 2018.

[13] W. Jin, R. Barzilay, and T. Jaakkola. Hierarchical generation of molecular graphs using structural motifs. In International Conference on Machine Learning, pages 4839–4848. PMLR, 2020.

[14] J. Jumper, R. Evans, A. Pritzel, T. Green, M. Figurnov, O. Ronneberger, K. Tunyasuvunakool, R. Bates, A. Žídek, A. Potapenko, et al. Highly accurate protein structure prediction with alphafold. Nature, 596(7873):583–589, 2021.

[15] I. Lee, J. Keum, and H. Nam. DeepConv-DTI: Prediction of drug-target interactions via deep learning with convolution on protein sequences. PLoS computational biology, 15(6):e1007129, 2019.

[16] T. Liu, Y. Lin, X. Wen, R. N. Jorissen, and M. K. Gilson. BindingDB: a web-accessible database of experimentally determined protein–ligand binding affinities. Nucleic acids research, 35(suppl_1):D198–D201, 2007.

[17] H. L. Morgan. The generation of a unique machine description for chemical structures-a technique developed at chemical abstracts service. Journal of Chemical Documentation, 5(2):107–113, 1965.

[18] B. Ramsundar. Molecular machine learning with DeepChem. PhD thesis, Stanford University, 2018.

[19] A. Rives, J. Meier, T. Sercu, S. Goyal, Z. Lin, J. Liu, D. Guo, M. Ott, C. L. Zitnick, J. Ma, et al. Biological structure and function emerge from scaling unsupervised learning to 250 million protein sequences. Proceedings of the National Academy of Sciences, 118(15), 2021.

[20] D. Rogers and M. Hahn. Extended-connectivity fingerprints. Journal of chemical information and modeling, 50(5):742–754, 2010.

[21] A. Shrikumar, P. Greenside, and A. Kundaje. Learning important features through propagating activation differences. In International Conference on Machine Learning, pages 3145–3153. PMLR, 2017.

[22] S. Sledzieski, R. Singh, L. Cowen, and B. Berger. D-SCRIPT translates genome to phenome with sequence-based, structure-aware, genome-scale predictions of protein-protein interactions. Cell Systems, 12:1–14, Oct. 2021.

[23] M. Tsubaki, K. Tomii, and J. Sese. Compound–protein interaction prediction with end-to-end learning of neural networks for graphs and sequences. Bioinformatics, 35(2):309–318, 2019.

[24] M. Zitnik, R. Sosic?, S. Maheshwari, and J. Leskovec. BioSNAP Datasets: Stanford biomedical network dataset collection. http://snap.stanford.educi U S A, Aug. 2018.

